# CA-170 – a potent small-molecule PD-L1 inhibitor or not?

**DOI:** 10.1101/662668

**Authors:** Bogdan Musielak, Justyna Kocik, Lukasz Skalniak, Katarzyna Magiera-Mularz, Dominik Sala, Mirosława Czub, Tad A. Holak, Jacek Plewka

## Abstract

**CA-170** is currently the only small-molecule modulator in clinical trials targeting PD-L1 and VISTA proteins – important negative checkpoint regulators of immune activation. The reported therapeutic results to some extent mimic those of FDA-approved monoclonal antibodies overcoming the limitations of the high production costs and adverse effects of the latter. However, no conclusive biophysical evidence proving the binding to hPD-L1 has ever been presented. Using well-known *in vitro* methods: NMR binding assay, HTRF and cell-based activation assays, we clearly show that there is no direct binding between **CA-170** and PD-L1. To strengthen our reasoning, we performed control experiments on **AUNP-12** – a 29-mer peptide, which is a precursor of **CA-170**. Positive controls consisted of the well-documented small-molecule PD-L1 inhibitors: **BMS-1166** and peptide p57.

## 1. Introduction

Alongside chemo- and radiotherapy, surgery, and other “targeted treatments”, cancer immunotherapy (called also immuno-oncology) is now regarded as the fifth pillar of cancer treatment, mainly due to a rapid development of potent immune checkpoint-blocking (ICB) therapeutic inhibitors [1–4]. These therapies unleash the native immune system by overcoming tumor-induced immunosuppression demonstrating impressive results. Among them, ICB agents, anti-PD-1/PD-L1 modulators have recently gained momentum and well-deserved acknowledgement in both academia (Nobel Prize in Physiology or Medicine in 2018 for James Allison and Tasuku Honjo for their discovery of cancer therapy by inhibition of negative immune regulation [5,6]) and pharmaceutical market ($5 billion in 2016 [7] and over 1500 different clinical studies on PD-1/PD-L1 agents as of 2017 (comprising mostly of combination therapies) [8]).

Programmed cell death protein 1 (known also as PD-1 and CD279) and its naturally occurring ligand PD-L1 (B7-H1, CD274) are transmembrane glycoprotein receptors characterized by β-sandwich immunoglobulin -like extracellular domains (seven β-strands organized in two sheets connected via a disulfide bridge) (Figure 1) [9,10]. PD-1 is expressed at the cell surface of activated T and B cells, monocytes, dendritic cells and natural killer (NK) T cells [11,12]. Its extracellular IgV domain is followed by a transmembrane region and an intracellular tail, which contains two tyrosine-based immunoreceptor signaling motifs: the switch motif (ITSM), and the inhibitory motif (ITIM) [13]. Similarly, human PD-L1 (hPD-L1) also contains extracellular IgV domain, responsible for the binding to human PD-1 (hPD-1), followed by the IgC domain and a transmembrane domain. The fully human PD-1/PD-L1 complex was first structurally characterized in 2015 [9] and is reported to have the 1:1 stoichiometry with partners interacting via the IgV domains by strands from the GFCC’ beta sheets of both proteins placing them perpendicular to each other. The interface has a total surface area of 1,970 Å^2^ and is maintained by polar and hydrophobic interactions. More polar interactions comprising, hydrogen bonds and salt bridges, are observed at the periphery of the hydrophobic core based mainly on alkyl-π and π-π interactions. Interestingly, only PD-1 undergoes significant structural rearrangement upon binding, most notably in the CC′ loop, which transits from an open conformation in the apo form to a closed one in C β strand [9].

**Figure 1.**
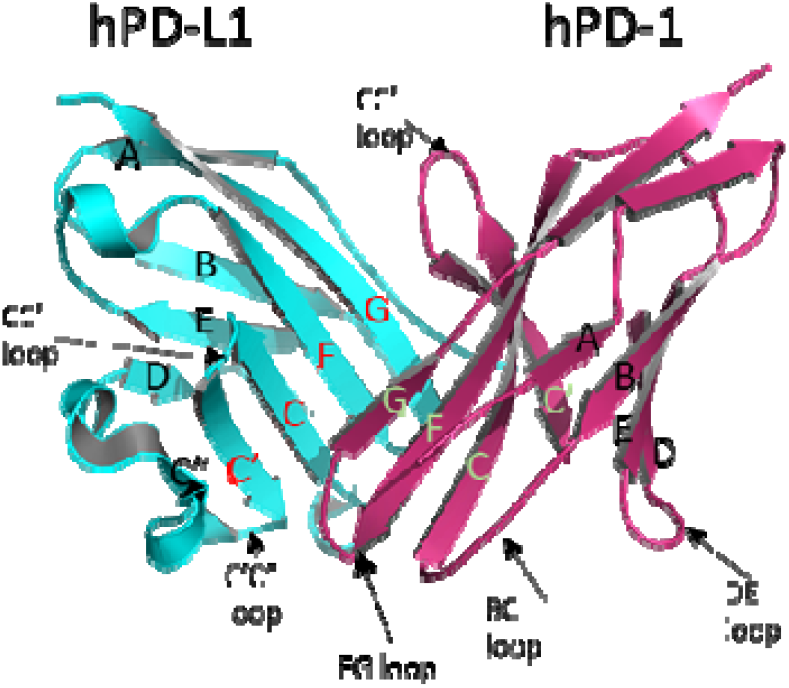
The structure of the hPD-1/hPD-L1 complex. hPD-L1 in cyan, hPD-1 in purple. Strands from IgV with names. Strands involved in protein-protein interaction (PPI) are coded in red for hPD-L1 and green for hPD-1 (adapted from [9]).

Physiologically, the PD-1/PD-L1 complex negatively regulates the immune response and provides so called T cell homeostasis [14,15]. Cancer cells often abuse this mechanism by overexpressing PD-L1 as a strategy to suppress anti-cancer activity of the immune system [16,17]. Therefore, disrupting the PD-1/PD-L1 complex at the cancer cell-T cell interface restores lymphocyte activity and has become an attractive target for pharmaceutical companies resulting in the development of potent monoclonal antibody-based therapies, that have demonstrated remarkable therapeutic successes. Immunotherapies based on monoclonal antibodies (mAbs) are limited, though, by the properties of relatively big protein agents; namely the need for intravenous administration, immune-related adverse effects or high production costs [18–20]. Small molecule-based therapeutics are believed to serve as a remedy for those shortcomings, in most cases being orally bioavailable and cheaper in manufacturing, while presenting improved pharmacokinetics and with greater diffusion rates [21,22]. However, as of today, there are no FDA-approved small-molecule modulators for the PD-1/PD-L1.

In the recent years, several attempts for the development of non-mAb PD-1/PD-L1 inhibitors, such as macrocyclic peptides, peptidomimetic molecules and nonpeptidic small-molecules, were presented and disclosed in particular by Bristol-Myers Squibb and Aurigene Discovery Technologies Limited (summarized in details in patent reviews [23–29]). In 2016, Aurigene in collaboration with Curis issued Phase I trials of **CA-170**, a small-molecule inhibitor against PD-L1/L2 and VISTA, with low nanomolar potencies for the treatment of advanced solid tumors and lymphomas (NCT02812875, clinicaltrials.gov). **CA-170** is also undergoing phase II clinical trials for lung cancer, head and neck / oral cavity cancer, MSI-H positive cancers and Hodgkin lymphoma in India (CTRI/2017/12/011026, ctri.nic.in). According to the preclinical *in vitro* and *in vivo* data reported by the company, **CA-170** demonstrated dose-dependent enhancement in the proliferation of PD-L1, PD-L2, and VISTA-inhibited T lymphocytes, and exhibited antitumor effects similar to those of antibodies, including tumor shrinkages and prolonged stable disease without adverse effects [30–34].

Aurigene and Curis have not disclosed the structures of **CA-170**, however, based on the analysis of Aurigene patents and the fact that it is derived from the **AUNP-12** peptide with known structure [35] CA 170 is most likel the compound 4 from the patent WO 2015/033301 A [36] being a peptidomimetic, composed of a serine, D-asparagine and threonine connected via diacylhydrazine and urea linker moieties, as presented in Figure 2a. The same structures are advertised under the name **CA-170** on following webpages of chemical vendors [37–40]. Its precursor, **AUNP-12** peptide (Figure 2b), was selected based on a rational structure-activity relationship (SAR) study, and presented a 100% activity in the mouse splenocyte proliferation assay [41].

**Figure 2.**
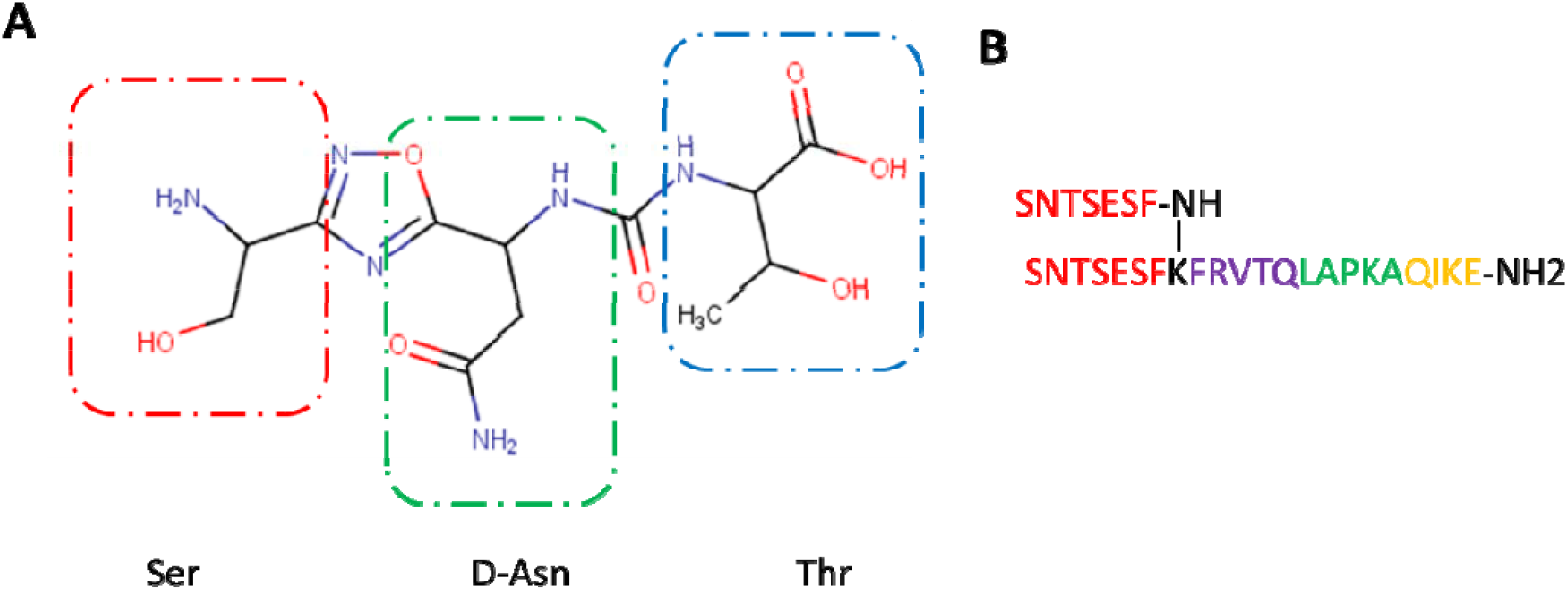
**A)** The putative structure of **CA-170** with building blocks indicated by boxes. **B)** Structure of the **AUNP-12** peptide composed of 4 hPD-1 parts: 2x BC loop (SNTSESF) in red connected via lysine to the D strand (sequence FRVTQ), purple; the FG loop (sequence LAPKA) in green, and the G strand (sequence QIKE), orange.

Despite being described as extremely potent with EC_50_ values of 17 and 0.72 nM for **CA-170** and **AUNP-12** peptide, respectively [36,41], no biophysical data providing direct binding to any of the reported targets is available no biophysical data providing direct binding to any of the reported targets is available, which has already rose some concerns suggesting that these compounds may act on the broader PD-1/PD-L1 pathway rather than the mentioned proteins itself [23]. Here, we employed a number of standard biophysical and biochemical methods, commonly used to assess the binding of tested molecules to the target proteins, in a conclusive way to determine affinities of **CA-170**, and its precursor **AUNP-12**, to hPD-L1. We also compare these molecules to the two well-characterized reference compounds – small molecule **BMS-1166** and the macrocyclic peptide p57 [42].

## 2. Results

### 2.1. **CA-170** does not bind to hPD-L1 according to the NMR binding assay

Nuclear magnetic resonance (NMR) is routinely employed to characterize small proteins, protein-protein and ligand-protein interactions [43]. Performing simple 2D experiment, such as heteronuclear multiple quantum coherence spectroscopy (HMQC) yields a two-dimensional map with a peak for each unique proton attached to heteroatom creating a kind of an individual fingerprint for each protein. Using this approach, one could verify if a given agent binds to the target protein by observing, whether the addition of tested agent induces changes of the initial protein spectrum [44]. The extent of those chemical shifts on the spectrum can be quantified to determine the affinity of tested molecule.

The affinities of **CA-170** and **AUNP-12** to hPD-L1 were assessed with the ^1^H and ^1^H–^15^N HMQC NMR spectroscopy. In the first experiment, the **CA-170** compound was titrated against the ^15^N-labeled hPD-1-binding single domain of hPD-L1 (hPD-L1(amino acids 18-134) and against the entire extracellular domain of the hPD-L1 protein (residues 18-239). In both 1D and 2D NMR spectra, no changes were observed for signals of the protein of hPD-L1, even when the 10-fold excesses of compound was used (Figure 3 a,b). These results indicate that **CA-170** does not bind to hPD-L1 even at the elevated concentration as no peak shift or broadening of the NMR signals were observed.

**Figure 3.**
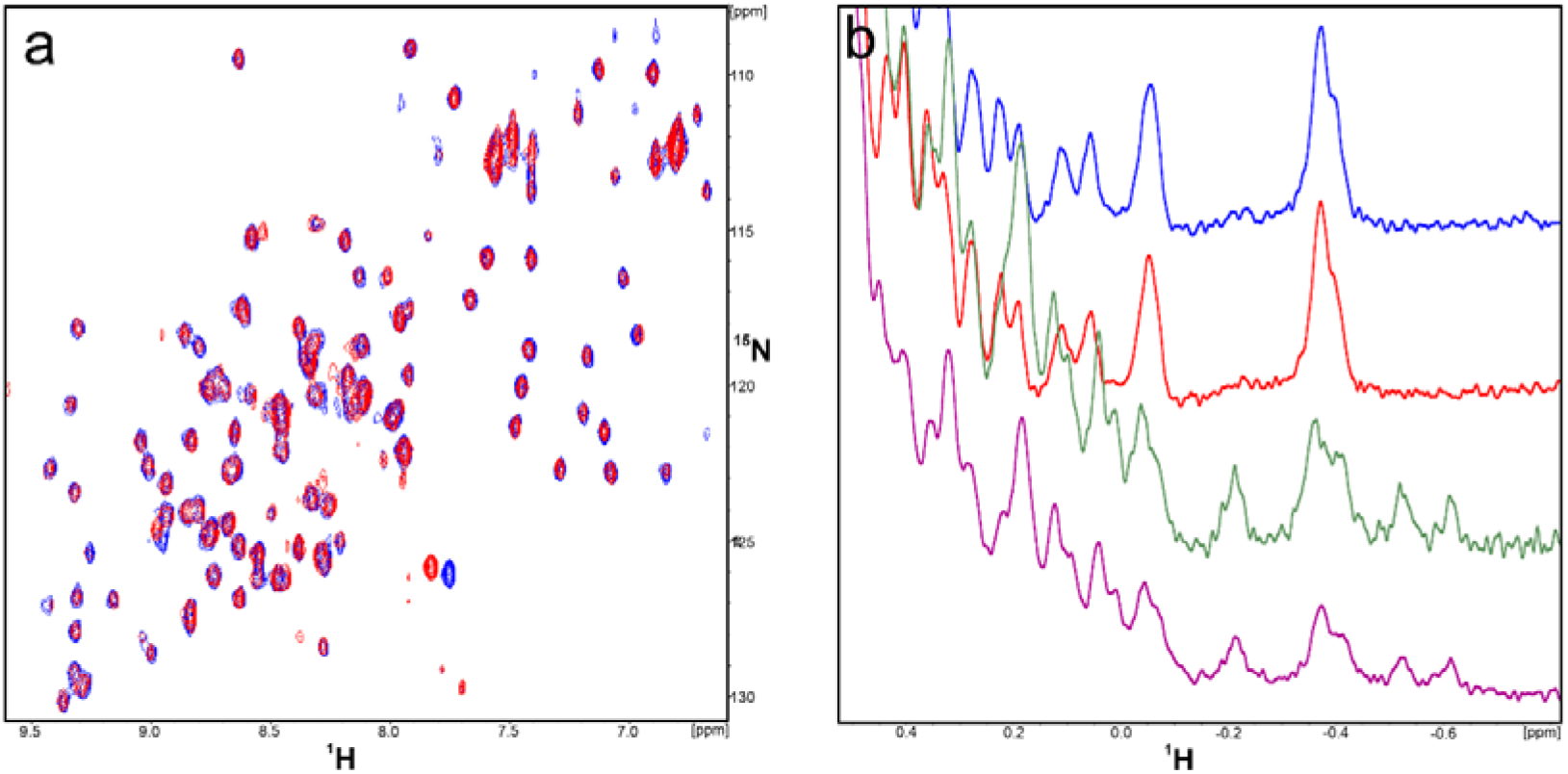
**A)** ^1^H-^15^N HMQC spectra of apo-hPD-L1(18-134) (blue) and hPD-L1(18-134) with **CA-170** (red) in the molar ratio protein/**CA-170** 1:10, respectively. **B)** ^1^H NMR spectra of apo-hPD-L1(18-134)(blue), hPD-L1(18-134) with **CA-170** (red) in the molar ration 1/10, apo-hPD-L1(18-239)(purple), and hPD-L1(18-239) with **CA-170** (green) in the molar ration 1/10. In none of spectra any interactions of the tested compound with the hPD-L1 protein were observed as the recorded peaks overlay each other.

Although **CA-170** is advertised as PD-L1 binder, we tested if it exhibits any interactions with hPD-1 to test all possibilities of disrupting the PD-1/PD-L1 complex. However, as in previous results, no interaction with hPD-1 protein was observed (Figure 4a,b). Since the selection method for the compound and *in vitro* assays were performed on the murine system showing spectacular results, we decided to conduct the same NMR binding experiment using mouse PD-L1 (mPD-L1) (Figure 2b). However, once again we did not observe any indication of the binding between the **CA-170** compound and mPD-L1.

**Figure 4.**
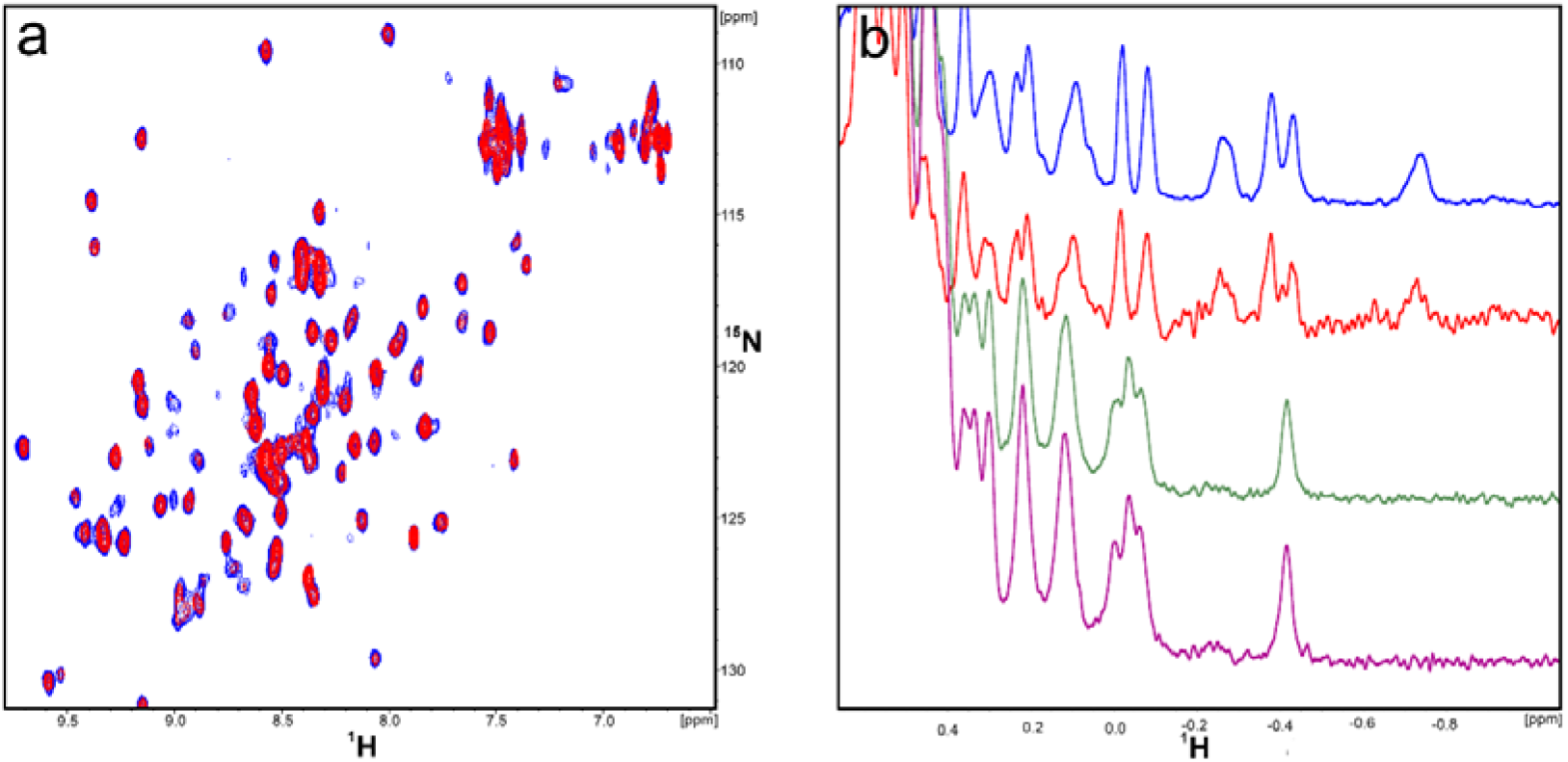
**A)** ^1^H-^15^N HMQC spectra of apo-hPD-1 (blue) and hPD-1 with **CA-170** (red) in the molar ration 1/10. **B)** ^1^H NMR spectra of apo- hPD-1 (blue), hPD-1 with **CA-170** (red) in the molar ratio 1/10, apo- mPD-L1 (green), and mPD-L1 with **CA-170** (purple) in the molar ratio 1/10.

Since **CA-170** was derived from the **AUNP-12** peptide, which is composed of the 4 fragments of hPD-1, we decided to also perform the NMR binding assay to test the affinity of **AUNP-12** to hPD-L1 in the same way. The analysis of perturbations of chemical shifts of protein NMR signals in the HMQC spectra of ^15^N hPD-L1 allowed us to determine that the **AUNP-12** binds to the short construct of hPD-L1(18-134) with the Kd at the millimolar level indicated by slight chemical shift perturbations (Figure 1aS). Unambiguous changes in the NMR spectra are observed for short hPD-L1 with the excess of **AUNP-12** at the molar ratio of 1/5. The rest of the results of NMR experiments for **AUNP-12** with hPD-L1(18-239) and with hPD-1 showed no interaction of the peptide and the proteins (Figures 1S b, 2S a,b).

Therefore, we determined in a conclusive way that neither **CA-170** nor **AUNP-12** binds to human or mouse PD-L1 or human PD-1. As the reference positive controls for the NMR binding assay, we show the NMR spectra of the well-known hPD-L1 binders: a macrocyclic peptide p57 developed by Bristol-Myers Squibb and the small molecule **BMS-1166**, also developed by Bristol-Myers Squibb, which bind strongly to hPD-L1 at equimolar concentrations (Figure 3S) (Adapted from Skalniak et al., Oncotarget, 2017 [42]).

### 2.2. **CA-170** cannot disrupt hPD-1/hPD-L1 complex as determined with HTRF assay

Regardless of the NMR measurements, we performed the IC_50_ determination for **CA-170** and **AUNP-12** peptide with the hPD-1/hPD-L1 complex using Homogenous Time Resolved FRET (HTRF), which is currently a standard methodology to determine potency of the inhibitors in numerous publications and patents, due to its robustness and absence of false-positive results [45]. We used increasing concertation of the inhibitors against standardized 5 nM of hPD-L1 and 50 nM of hPD-1 in duplicates. The resulting normalized data was then fitted with the Hill’s equation. We also determined IC_50_ of well-known hPD-L1 inhibitors as the positive reference: **BMS-1166** and macrocyclic peptide p57 to cross-validate the results with the literature (Figure 5).

**Figure 5.**
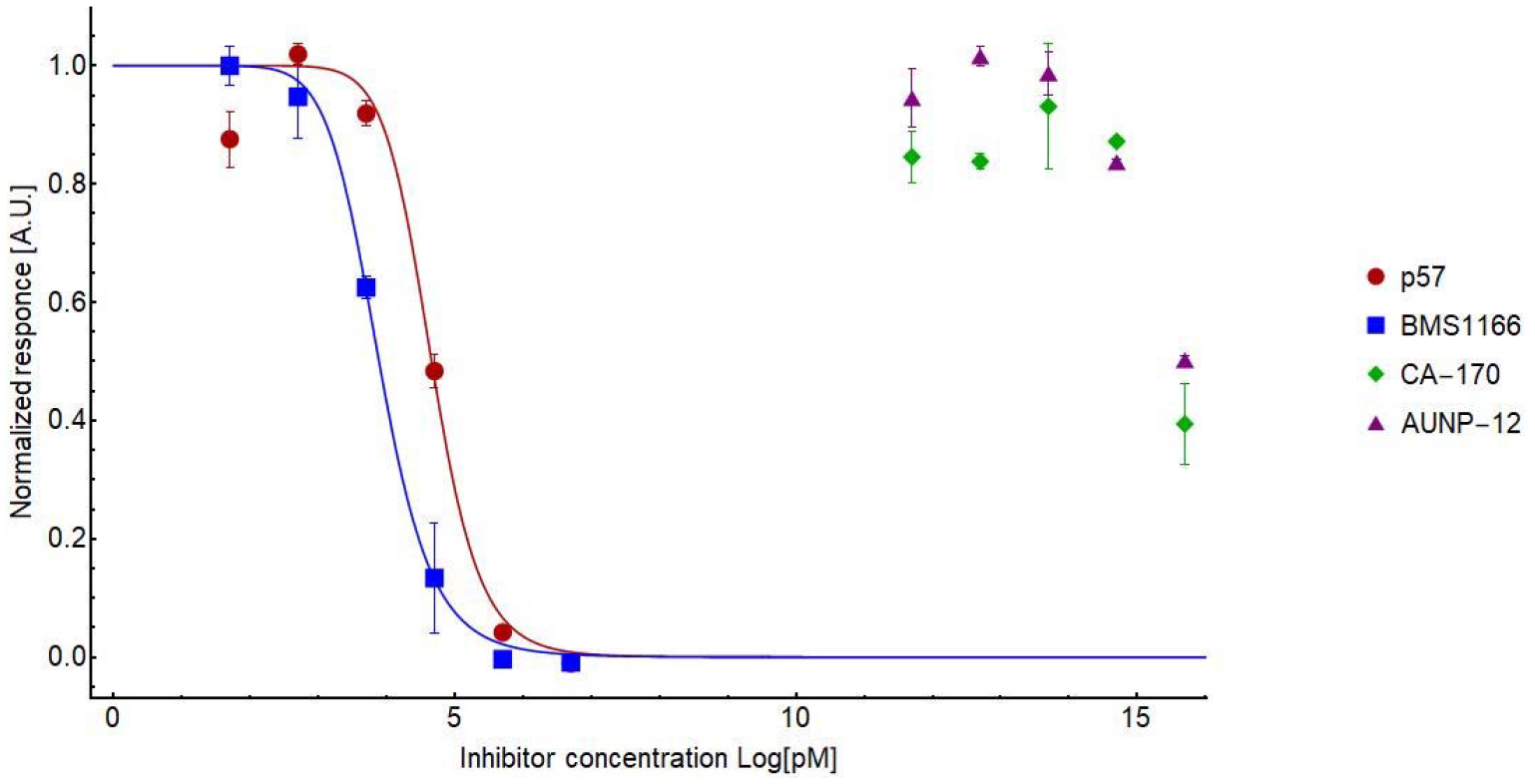
Determination of the potency for reference compounds peptide p57 (red circles), **BMS-1166** (blue squares) as compared to almost no response from **CA-170** (green diamonds), and **AUNP-12** (violet triangles). Fitting was performed using normalized Hill’s equation.

**Figure 6.**
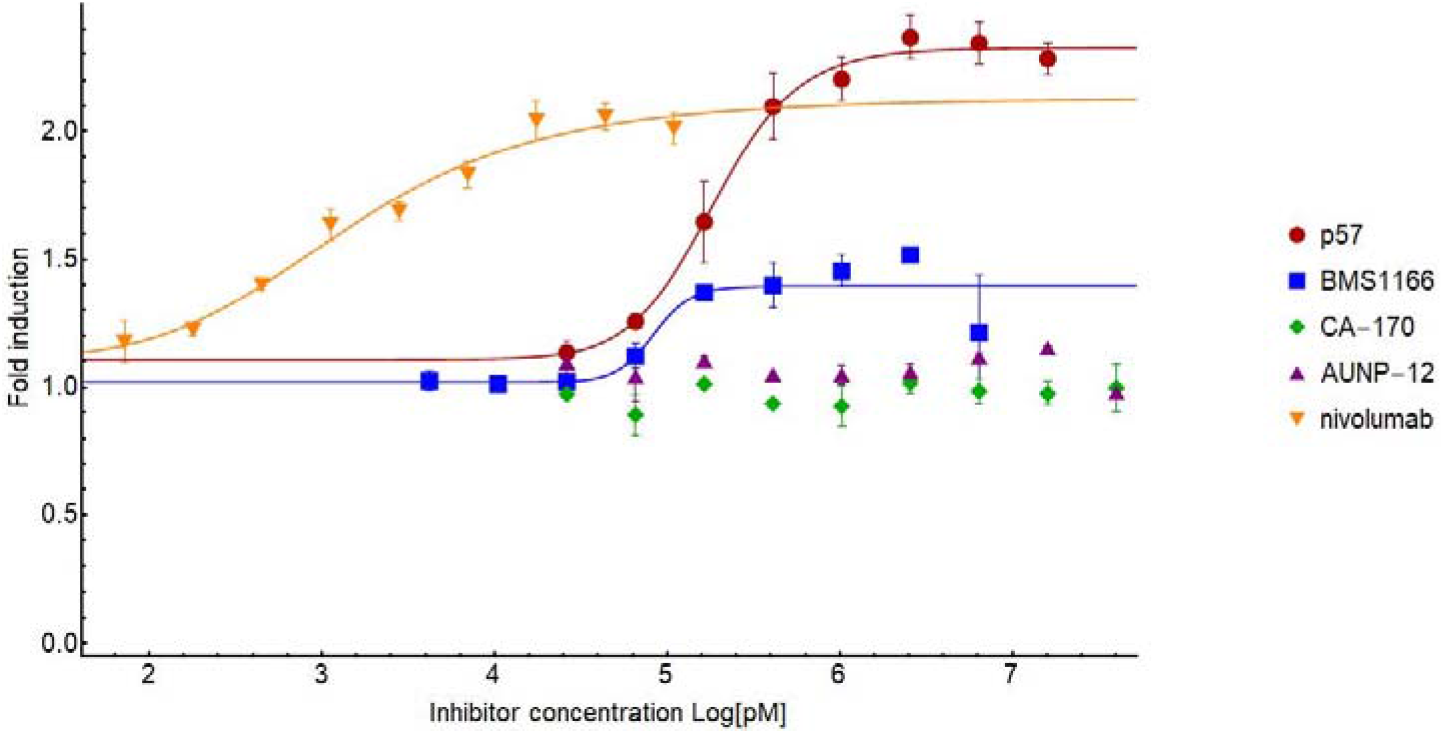
The comparison of bioactivities of nivolumab (orange triangles), peptide p57 (red circles), **BMS-1166** (blue squares), **CA-170** (green diamonds), and **AUNP-12** (violet triangles) in the hPD-1/hPD-L1 immune checkpoint assay. Fitting was performed using a normalized Hill’s equation.

From the above consideration, it is clear that both reference compounds are potent in dissociating the hPD-1/hPD-L1 complex with IC_50_ of 45.4+/− 0.001 nM and 7.7+/−0.01 nM for peptide p57 and **BMS-1166**, respectively, which is in hand with the values reported in the literature [22,42]. However, both **AUNP-12** and **CA-170** only marginally dissociated the complex at the highest used concentration of 5 mM indicating that the estimated IC_50_ would be above 5-10 mM. It should be noted, though, that at such high concentrations the interactions of small molecules with isolated protein complex may be unspecific. Therefore, we presented in a conclusive way that using standard biophysical methods for assessing affinities of a small compound towards target proteins by NMR and HTRF assays, neither **AUNP-12** nor **CA-170** exhibits potency in dissociation of the hPD-1/hPD-L1 complex.

### 2.3. **CA-170** fails to restore the activation of hPD-1/hPD-L1-blocked effector Jurkat T cells

To verify the activity of compound **CA-170** in a hPD-1/hPD-L1-sensitive cell line test, the compound was challenged in the immune checkpoint blockade bioassay [45]. For this, antigen-presenting CHO-K1 cells overexpressing hPD-L1 and T Cell Receptor (TCR) Activator (hPD-L1 aAPCs) were co-cultured with the effector Jurkat T cells overexpressing hPD-1 and a luciferase gene controlled by the NFAT-Response Element (hPD-1 Effector Cells, hPD-1 ECs). In the assay, the activation of TCR-signaling by the TCR-activator is inhibited by the hPD-1/hPD-L1 interaction. Upon hPD-1/hPD-L1 blockade, normal Jurkat T cell activation is restored, as monitored by the increased luciferase activity.

In the immune checkpoint blockade assay, three positive control molecules were used: small molecule **BMS-1166**, a macrocyclic peptide p57, and an approved monoclonal antibody nivolumab [47]. All three compounds restored the activation of hPD-1 ECs, however significant differences in their potencies were noticed (Figure 7). According to the assay, the EC50 for nivolumab was 1.4 nM, for **BMS-1166** – 83.4 nM, and for peptide p57 - 185.5 nM, which are close to the values reported in the HTRF assay. This observation is also in agreement with our previous reports, where we have shown that the macrocyclic peptides and BMS compounds, although active, present much higher EC_50_ values and lower maximal effects compared to therapeutic antibodies [42,46]. On the other hand, neither **CA-170**, nor its precursor **AUNP-12**, was able to provide the activation of Jurkat T cells, repressed by the hPD-1/hPD-L1 checkpoint, up to the concentration of 40 μM of the compounds. This is a strong evidence that the compounds fail to interfere specifically with the hPD-1/hPD-L1 immune checkpoint in the biological system.

**Figure 7.**
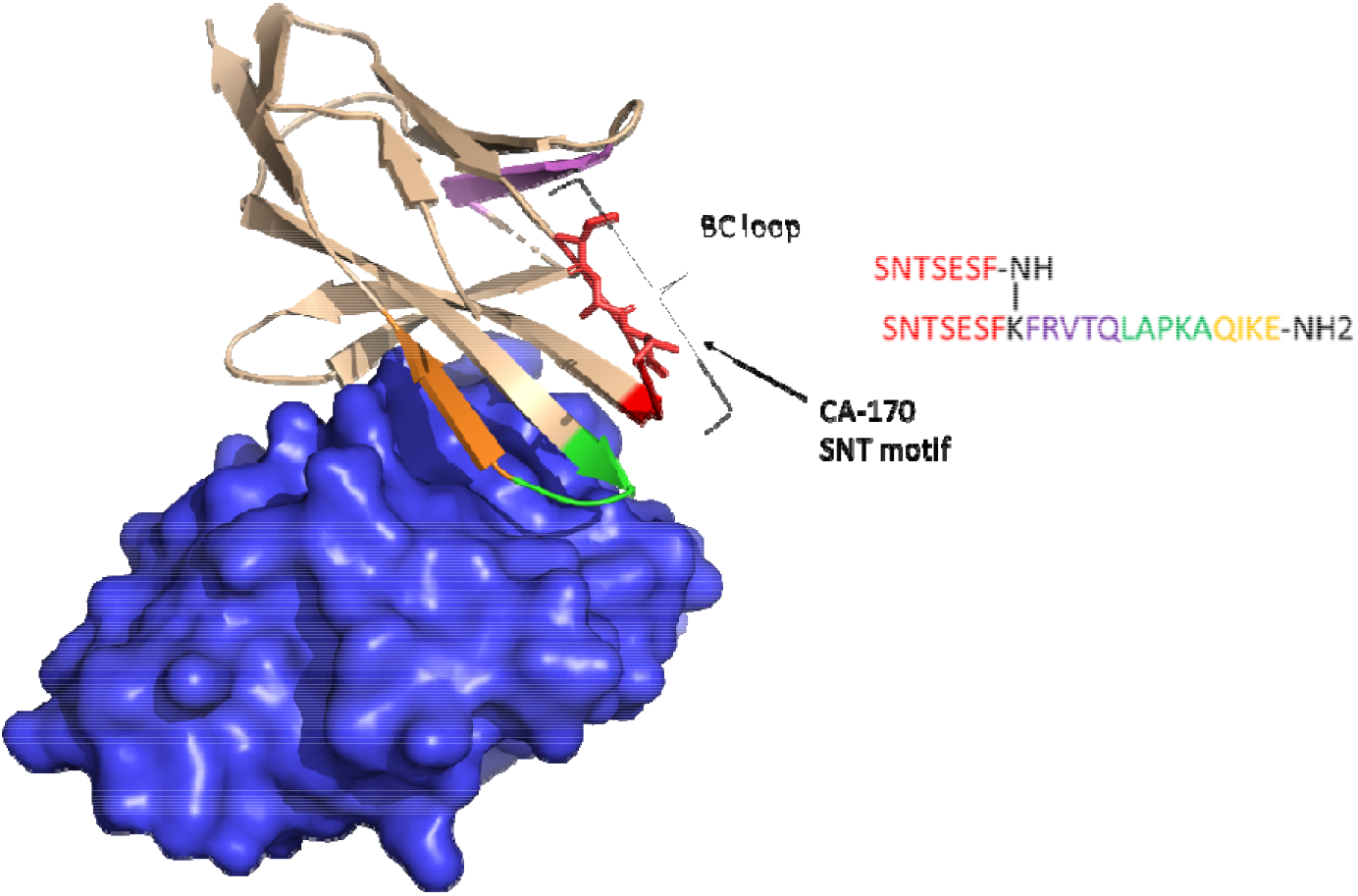
hPD-1 (light brown cartoon) and hPD-L1 (blue surface) in complex from 4ZQK PDB. The BC loop is highlighted. The **AUNP-12** peptide sequence (on the right) is color coded on corresponding strands of hPD-1. SNT-motif from **CA-170** is represented as sticks in BC loop.

## 3. Discussion

Small molecule modulators for the hPD-1/hPD-L1 pathway are of a great interest to both patients and pharmaceutical companies. This is reflected by the increasing number of patent applications for small-molecular scaffolds. HTRF or AlphaScreen assay are used as standardized methods to determine the potency of screen compounds in disrupting the hPD-1/hPD-L1 complex. However, even though many of them exhibits sub-nanomolar affinities, none of them progressed to the clinical trials, due to multiple reasons, such as poor drugability, toxicity profile of the compounds, etc. The only compound currently in clinical trial phase I is **CA-170**, which is advertised as the modulator that dually targets VISTA and hPD-L1. The structure of **CA-170** has not been formally disclosed. Based on the patent and review analysis, we have concluded that it is the compound 4 from the patent WO 2015/033301 A [36]),. Moreover, Aurigene/Curis did not provide any biophysical results confirming direct binding of their compound to any of the advertised targets, neither for **CA-170** nor to its precursor **AUNP-12**, raising some speculations regarding the mechanism of the action of these modulators. The only confirmation of the potency of **CA-170** and **AUNP-12** comes from the proliferation restoration assays and IFN-γ secretion rescue from mouse splenocyte and human PBMC. Lack of orthogonal methods for the confirmation of the potency and inconclusive nature of applied methodology made us question the mechanism of action for **CA-170** and **AUNP-12**. Therefore, we conducted our own investigation to determine the binding affinities of **CA-170** and **AUNP-12** to hPD-L1 in a conclusive way using well acknowledged methods of NMR binding assay, HTRF and the hPD-1/hPD-L1 immune checkpoint blocking assay.

Based on the above consideration we state that neither **CA-170** nor its precursor **AUNP-12** exhibit any binding to hPD-L1 that would be strong enough to disrupt the hPD-1/hPD-L1 complex. Furthermore, they do not bind to mouse PD-L1 (mPD-L1) or hPD-1 either. We, however, do not exclude the possibility that these compounds may act downstream from the hPD-1 receptor or on any other T cell-activating pathway, since the compounds clearly present promising therapeutic results.

We hypothesize that the reason why Aurigene compounds do not bind to hPD-L1 may be explained by the conceptual fault in the initial “rational design”. A design starting point, the compound **AUNP-12**, is a hybrid of the peptide sequences extracted from hPD-1, that are supposed to be close to its interaction surface with hPD-L1. However, if we look at the locations of particular peptide fragments of **AUNP-12**. (Figure 8), it shows that only the LAPKAQIKE sequence is present at the direct interface of the hPD-1/hPD-L1 interaction. This might be the reason why **AUNP-12** presents some, albeit weak, binding to hPD-L1 in the NMR assay at high compound concentrations. However, the BC loop containing the SNT motif, that constitutes the optimized **CA-170** compound, is pointing away from the PD-1/PD-L1 interface as can be inferred from the fully human PD-1/PD-L1 complex structure first reported by Zak et. al in 2015 [9]. This explains the lack of direct binding between hPD-L1 and **CA-170**, which does not justify the rationality of the selection of this particular part of hPD-1 as a putative inhibitor.

Therefore, both **AUNP-12** and **CA-170** are not targeted at hPD-L1 as advertised, which may still be in agreement with the results from the splenocyte screening assay, given that the assay does not have to be hPD-1/hPD-L1 specific due to its complex nature. Despite above consideration, **CA-170** shows promising results in clinical trials and further investigation of the mode of action, especially identifying the binding partner/s, is highly recommended.

## 4. Materials and Methods

### 4.1. Materials

**CA-170** was purchased from MedChemExpress Cas: 1673534-76-3, **AUNP-12** was purchased from SelleckChem, **BMS-1166** was synthesized according to the protocol from described in [22], peptide p57 was synthesized as described in [48].

### 4.2. Protein expression and purification

The gene encoding hPD-1 (amino acids 33 – 150, Cysteine 93 mutated to serine) was cloned in to pET-24d, the gene encoding hPD-L1 (amino acids 18 - 134 with C-terminal HisTag) was cloned into pET-21b, the genes encoding hPD-L1 (amino acids 18 - 239 with C-terminal HisTag) and mPD-L1 (amino acids 38 – 134) was cloned into pET-28a respectively. Proteins were expressed in the *Escherichia coli* BL21 (DE3). Bacterial cells were cultured in LB or M9 minimal medium containing ^15^NH_4_Cl as a nitrogen source in ^15^N labeling at 37°C. Proteins expression was induced with 1 mM Isopropyl β-D-1-thiogalactopyranosid (IPTG) at OD_600_ of 0.8 and the cells were cultured overnight. For hPD-1, hPD-L1 and mPD-L1 after induction temperature was lowered to 28°C, for hPD-L1(18-239) temperature was stabled at 37°C. Inclusion bodies purification was carried out as described previously [9]. Afterwards inclusion bodies purification proteins were refolded by drop-wise dilution into solution containing 0.1 M Tris pH 8.0, 0.4M L-Arginine hydrochloride,), 2 mM EDTA, 5 mM cystamine and 0.5 mM cysteamine for hPD-1 and 0.1 M Tris pH 8.0 containing 1 M L-Arg hydrochloride, 0.25 mM oxidized glutathione and 0.25 mM reduced glutatione for hPD-L1, human PD-L1(18-239) and mPD-L1 respectively. After refolding proteins were dialyzed 3 times against solution containing 10 mM Tris pH 8.0 and 20 mM NaCl. Later, proteins were purified by SEC (size-exclusion chromatography) on HiLoad 26/600 Superdex 75 column (GE Healthcare) in 25 mM sodium phosphate pH 6.4 with 100 mM NaCl for hPD-1 or in PBS pH 7.4 for hPD-L1, hPD-L1(18-239) and mPD-L1 respectively.

### 4.3. NMR binding assay

For NMR measurements, the buffer was exchanged by gel filtration to PBS pH 7.4. 10% (v/v) of D_2_O was added to the samples to provide the lock signal. All spectra were recorded at 300 K using a Bruker Avance III 600 MHz spectrometer. Binding of the compounds was analyzed by titrating the ^15^N-labeled hPD-L1/hPD-1 and recording the ^1^H and ^1^H−^15^N HMQC spectra prior to and after the addition of the compounds.

### 4.4. Homogenous Time Resolved FRET

HTRF assay was performed using the certified Cis-Bio assay kit at 20 μL final volume using their standard protocol (5 nM of h-L1 and 50 nM of hPD-1 in the final formulation). To determine the half maximal inhibitory concentration (IC_50_) of tested compounds, measurements were performed on individual dilution series. After mixing all components according to Cis-Bio protocol, the plate was left for 2h incubation at room temperature followed by TR-FRET measurement on Tecan Spark 20M. Collected data was background subtracted on the negative control, normalized on the positive control, averaged and fitted with normalized Hill’s equation to determine the IC_50_ value using Mathematica 12.

### 4.5. Cell culture

CHO K-1 cells overexpressing hPD-L1 and the recombinant TCR ligand (hPD-L1 Antigen Presenting Cells, hPD-L1 aAPCs, Promega) and Jurkat T cells overexpressing hPD-1 and carrying a luciferase reporter gene under the control of Nuclear Factor of Activated T-cells Response Element (NFAT-RE) (hPD-1 Effector Cells, hPD-1 ECs, Promega) were cultured in RPMI-1640 medium (Biowest) supplemented with 10% Fetal Bovine Serum (FBS, Biowest) and 200 mM L-Glutamine (Biowest) in the presence of G418 (250 μg/ml, InvivoGen) and Hygromycin B Gold (50 μg/ml, InvivoGen) as selection antibiotics. The overexpression of PD-L1 and TCR ligand in aAPCs and PD-1 in ECs were confirmed by flow cytometry and western blot analysis, respectively. PCR tests for Mycoplasma sp. contamination [48] were routinely performed and indicated negative results for both cell lines.

### 4.6. hPD-1/hPD-L1 immune checkpoint blockade assay

The activity of the inhibitors of hPD-1/hPD-L1 immune checkpoint was examined using the hPD-1/hPD-L1 Blockade Bioassay (Promega), according to the manufacturer’s instructions. hPD-L1 aAPCs were seeded on 96-well (white) plates at the density 10 000 cells/well 17 h prior to the experiment. The 2.5-fold dilutions of the small molecules or peptide 57 were first prepared in DMSO. On the day of the assay the compounds were diluted 1000-fold in the assay buffer (99% RPMI 1640, 1% FBS) to maintain the constant concentration of DMSO (0.1% of total volume). The 2.5-fold dilutions of nivolumab, a positive control anti-hPD-1 monoclonal antibody (Opdivo, Bristol-Myers Squibb), were prepared in the assay buffer on the day of the assay. The culture medium was discarded from the wells and serial dilutions of either the small-molecule or antibody was added. Afterwards, Jurkat hPD-1 cells were seeded at the density of 20 000 cells per well in the assay’s plates. After 6 h of the incubation in standard culture conditions, assay plates were equilibrated at ambient temperature for 10 min, followed by a 20 min incubation with the Bio-GloTM Assay reagent (Promega). The luminescence was detected using the Infinite M200 reader. Half maximal effective concentrations (EC_50_ values) were calculated from the Hill’s curve fitting to the experimental data.

## Supplementary Materials

The following are available online at xxx, Figure 1S A) ^1^H-^15^N HMQC spectra of apo-hPD-L1(18-134) (blue) and hPD-L1(18-134) with **AUNP-12** (red) in the molar ration 1/5. B) 1H NMR spectra of apo-hPD-L1(18-134)(blue), hPD-L1(18-134) with **AUNP-12** (red) in the molar ration 1/5, apo-hPD-L1(18-239)(purple), and hPD-L1(18-239) with **AUNP-12** (green) in the molar ration 1/5, Figure 2S. A) ^1^H-^15^N HMQC spectra of apo-hPD-1 (blue) and hPD-1 with **AUNP-12** (red) in the molar ration 1/5. B) ^1^H NMR spectra of apo- hPD-1 (blue), hPD-1 with **AUNP-12** (red) in the molar ration 1/5., Figure 3S. ^1^H NMR spectra of apo- hPD-L1(18-134) (blue) and with **BMS-1166** compound (green) and peptide p57 (red) in molar ration 1/1, respectively.

## Author Contributions

J.P. wrote the draft of the manuscript. All authors discussed the experiments and commented on the manuscript. B.M. performed NMR experiments; J.P and M. Cz. performed HTRF; J.S. and J.K. carried out the cell-based assays; K.M.-M. and D.S. provided support with preparation of expression plasmids and with protein purification.

## Acknowledgments

J.K. acknowledges the support of InterDokMed project no. POWR.03.02.00-00-I013/16

