## Supplementary Materials for "CA-170 – a potent small-molecule PD-L1 inhibitor or not?"

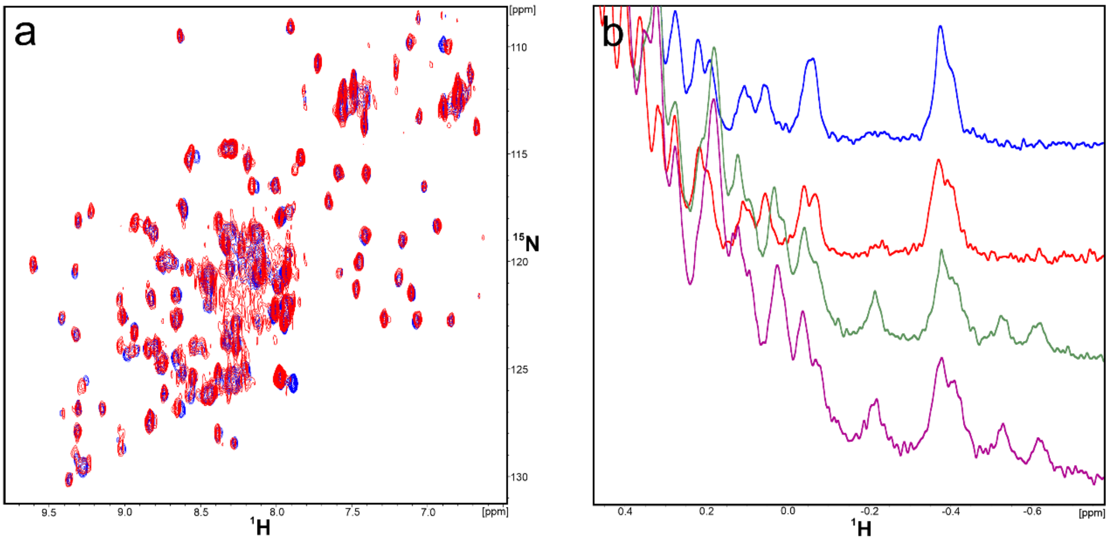


**Figure 1S. A)** $^{1}$H-$^{15}$N HMQC spectra of apo-hPD-L1(18-134) (blue) and hPD-L1(18-134) with AUNP-12 (red) in the molar ration 1/5. **B)** $^{1}$H NMR spectra of apo-hPD-L1(18-134)(blue), hPD-L1(18-134) with AUNP-12 (red) in the molar ration 1/5, apo-hPD-L1(18-239)(purple), and hPD-L1(18-239) with AUNP-12 (green) in the molar ration 1/5.

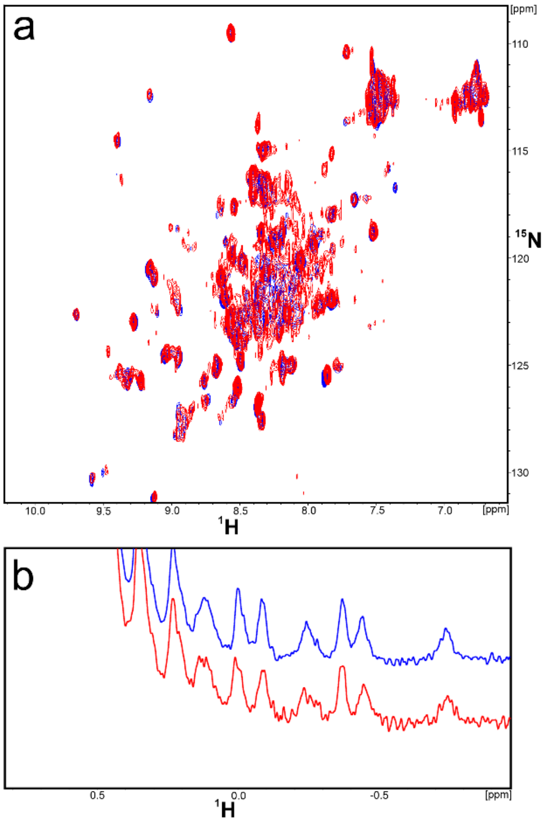


**Figure 2S. A)** $^{1}$H-15N HMQC spectra of apo-hPD-1 (blue) and hPD-1 with AUNP-12 (red) in the molar ration 1/5. **B)** $^{1}$H NMR spectra of apo- hPD-1 (blue), hPD-1 with AUNP-12 (red) in the molar ration 1/5.

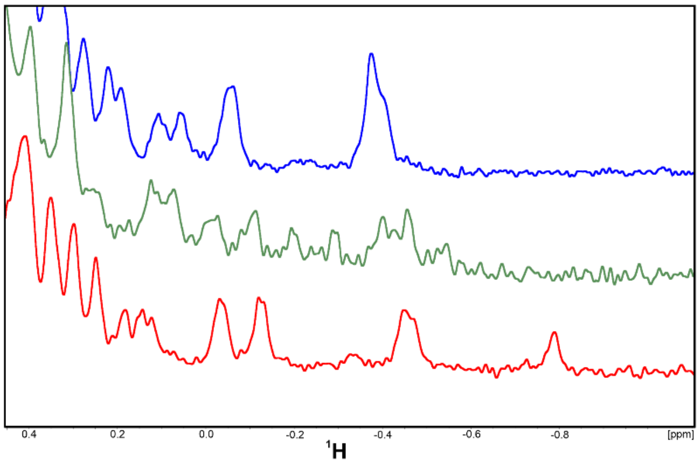


**Figure 3S.** $^{1}H$ NMR spectra of apo- hPD-L1(18-134) (blue) and with BMS-1166 compound (green) and peptide p57 (red) in molar ration 1/1, respectively.
